# The Effects of Posture on the Three-Dimensional Gait Mechanics of Human Walking in Comparison to Bipedal Chimpanzees

**DOI:** 10.1101/2021.07.30.454517

**Authors:** Russell T. Johnson, Matthew C. O’Neill, Brian R. Umberger

## Abstract

Humans walk with an upright posture on extended limbs during stance and with a double-peaked vertical ground reaction force. Our closest living relatives, chimpanzees, are facultative bipeds that walk with a crouched posture on flexed, abducted hind limbs and with a single-peaked vertical ground reaction force. Differences in human and bipedal chimpanzee three-dimensional kinematics have been well quantified; however, it is unclear what the independent effects of using a crouched posture are on three-dimensional gait mechanics for humans, and how they compare with chimpanzees. Understanding the relationships between posture and gait mechanics, with known differences in morphology between species, can help researchers better interpret the effects of trait evolution on bipedal walking. We quantified pelvis and lower limb three-dimensional kinematics and ground reaction forces as humans adopted a series of upright and crouched postures and compared them with data from bipedal chimpanzee walking. Human crouched posture gait mechanics were more similar to bipedal chimpanzee gait than normal human walking, especially in sagittal plane hip and knee angles. However, there were persistent differences between species, as humans walked with less transverse plane pelvis rotation, less hip abduction, and greater peak horizontal ground reaction force in late stance than chimpanzees. Our results suggest that human crouched posture walking reproduces only a small subset of the characteristics of three-dimensional kinematics and ground reaction forces of chimpanzee walking, with the remaining differences likely due in large part to the distinct musculoskeletal morphologies of humans and chimpanzees.

**Summary Statement:** Differences between human crouched posture gait and bipedal chimpanzee gait illustrate the limitations of using modern day humans to infer the evolution of hominin bipedalism.

## Introduction

Humans are unique among primates in the habitual use of an upright bipedal walking stride (Alexander, 2004). In general, the gait kinematics and ground reaction forces (GRFs) associated with our body posture during walking include adducted, extended lower limbs with little pelvis rotation, a vertically-oriented trunk and a double-peaked (i.e., bi-phasic) vertical GRF (Kadaba et al., 1989; O’Neill et al., 2015; Rose and Gamble, 2005). The gait mechanics exhibited by humans are distinct from that of common chimpanzees (*Pan troglodytes* (Blumenbach, 1775)), who are our closest-living relatives (Goodman, 1999; Waterson et al., 2005), and are facultative bipeds (e.g., Doran, 1992; Hunt, 1992; Sarringhaus et al., 2014). Bipedal kinematics and GRFs in chimpanzees are characterized by a posture that includes abducted, flexed hind limbs with greater rotations at the pelvis, an anteriorly-tilted trunk (Elftman, 1944; Jenkins, 1972; O’Neill et al., 2015) and a single-peaked (i.e., mono-phasic) vertical GRF (Kimura et al., 1977; O’Neill et al., in review; Pontzer et al., 2014). There is a longstanding interest in the extent to which the musculoskeletal morphology of the lower back, pelvis and lower/hind limbs underlie these differences in gait mechanics, with implications for understanding the evolution of bipedalism in hominins (e.g., DeSilva et al., 2018; Lovejoy, 2005; Lovejoy et al., 2009; Stern Jr. and Susman, 1983; Ward, 2002). A variety of approaches have been used to try to understand the relationship between morphological structure and gait mechanics, including making inferences about the gait mechanics of fossil hominins based on data from humans walking with crouched postures (Carey and Crompton, 2005; Crompton et al., 1998; Foster et al., 2013; Li et al., 1996; Raichlen et al., 2010; Wang et al., 2003; Yamazaki et al., 1979).

For this study, *posture* is defined as the general orientation of the body during walking (e.g., upright or crouched), while the term *gait mechanics* refers to specific variables (e.g., joint angles or GRFs) that vary throughout the stride. Previous studies of human crouched posture gait have sought to emulate bipedal chimpanzee walking (Li et al., 1996; Yamazaki et al., 1979) and/or sought to test hypotheses about walking in fossil hominins (Carey and Crompton, 2005; Crompton et al., 1998; Foster et al., 2013; Raichlen et al., 2010). These studies have focused solely on manipulating and measuring sagittal plane kinematics and kinetics. Yet, adopting broad changes in body posture during walking should be expected to affect gait mechanics outside of the sagittal plane. Humans can be provided instructions to walk with a crouched posture that imitates sagittal plane chimpanzee gait kinematics (Foster et al., 2013), but it is unknown whether the non-sagittal plane gait kinematics or GRFs for crouched walking in humans are similar to bipedal chimpanzees. The differences in musculoskeletal morphology between humans and chimpanzees could lead to differences in the overall three-dimensional (3-D) gait mechanics of crouched human walking and chimpanzee bipedal walking, which would imply a key limitation of manipulating posture in humans to infer how extinct hominins, with distinct musculoskeletal morphologies, would have walked.

In addition to a lack of quantitative data on the 3-D nature of crouched posture walking in humans, no studies focused on evolution of bipedalism have altered trunk orientation during crouched limb walking. However, trunk flexion represents a salient difference between human and bipedal chimpanzee gait (O’Neill et al., 2015; Pontzer et al., 2014; Thompson et al., 2015). The anterior trunk flexion in a bipedal chimpanzee shifts the whole-body center of mass forward, which may contribute to their flexed limb posture to place their feet under their center of mass (Lovejoy, 2005; Lovejoy and McCollum, 2010). Anterior trunk flexion in humans has been shown to affect lower limb kinematics and vertical GRF patterns (Aminiaghdam et al., 2017; Grasso et al., 2000; Kluger et al., 2014; Saha et al., 2008), although the most substantial effects require more extreme anterior trunk flexion (i.e. 50-90° relative to vertical) than observed in either bipedal chimpanzees (i.e., approx. 30-35°, Pontzer et al., 2014) or in other primates walking bipedally, such as macaques (i.e., approx. 25°, Ogihara et al., 2010). As such, it remains unclear how effects of moderate anterior trunk flexion characteristic of bipedal walking in non-human primates affects gait mechanics in humans. Further, the combined effects of trunk flexion and flexed limb posture on 3-D human gait kinematics and GRFs is unknown.

Along with the documented flexed limb posture during chimpanzee bipedal walking, there is also substantial non-sagittal plane motion that distinguishes it from normal human walking. Chimpanzees exhibit greater range of motion (ROM) than humans for some non-sagittal plane kinematic variables when walking at matched dimensionless speeds (O’Neill et al., 2015), yet perhaps the most striking difference is that pelvis list for chimpanzees is completely out-of-phase with human walking. In the single-support period, bipedal chimpanzees elevate their pelvis on the swing limb side, whereas humans drop their pelvis on the swing limb side (Jenkins, 1972; Kikel et al., 2020; O’Neill et al., 2015). Despite the opposite phasing of pelvis motion, this feature has not been included in human crouched posture studies. One study instructed humans to walk with an upright posture and a chimpanzee-like pelvis list motion, which resulted in greater hip abduction angles during stance than for normal human walking (Kikel et al., 2020). But it is unclear whether having humans emulate chimpanzee frontal plane pelvis motion during crouched posture walking will result in 3-D hip joint kinematics or GRF patterns similar to those observed in chimpanzees. However, even with multiple instructions given to humans to emulate chimpanzee gait mechanics, it is likely that some differences will persist given morphological differences between species.

Human lower limb, pelvis, and trunk posture can be easily manipulated during gait by providing instructions and feedback to participants. Human crouched posture instructions allow us to experimentally manipulate posture, while controlling for other musculoskeletal and physiological traits. As such, human crouched posture gait studies have the potential to provide important insights into how different musculoskeletal factors contribute to gait mechanics. However, a full 3-D accounting of differences in gait kinematic and GRFs across postures is required, as crouched bipedal walking may have substantial motion and forces outside of the sagittal plane, as exhibited by chimpanzees (O’Neill et al., 2015; O’Neill et al., in review). Quantifying these differences will lead to a greater understanding of the effect of posture on gait mechanics and allow for inferences about how the morphology of humans and chimpanzees give rise to their distinct gait patterns. Here, we instructed humans to walk with a series of crouched postures to evaluate how instructions focused on lower limb flexion, anterior trunk flexion, and pelvis motion lead to alterations in the 3-D pelvis and lower limb kinematics and GRFs. Specifically, we collected 3-D gait data for (i) normal human walking, (ii) walking with a crouched-limb, (iii) walking with a crouched-limb and anteriorly flexed trunk, and (iv) walking with a crouched-limb, anteriorly flexed trunk, and a chimpanzee-like pelvis list pattern. The normal and three crouched posture conditions for humans were compared with previously collected bipedal chimpanzees data to determine the degree to which crouched posture human walking is similar to that of chimpanzees (O’Neill et al., 2015; O’Neill et al., in review). We hypothesized that the normal human walking condition would be the least similar to chimpanzee gait mechanics out of all human conditions. We further hypothesized that the CL, CLT, and CLTP conditions would lead to progressively more chimpanzee-like gait mechanics.

## Materials and Methods

### Human Protocol

Ten healthy human subjects (5 female/5 male; age: 27 ± 5 years; height: 1.70 ± 0.08 m; mass: 68 ± 11 kg; lower limb length: 0.88 ± 0.04 m) were recruited for this study. All subjects had no history of gait pathologies, cardiovascular disease, neurological disease, or orthopedic problems that would affect how they walked. Due to the requirement to complete numerous walking trials with several different postures, we recruited subjects who were at least recreationally active to reduce the risk of fatigue being a factor during our data collection. All subjects self-reported that they met the American College of Sports Medicine recommendation of at least 150 minutes of moderate-to-vigorous physical activity per week (Garber et al., 2011). Prior to participating, subjects read and signed an informed consent document approved by the University of Massachusetts Amherst institutional review board. Subjects also completed a Physical Activity Readiness Questionnaire to assess their readiness to participate in the study (Thomas et al., 1992; Warburton et al., 2011).

For each subject, height, mass, age, and lower limb length were recorded. A set of retro-reflective markers were attached to anatomical landmarks on the upper body, pelvis, and lower limbs, and clusters of non-colinear markers were placed on the thigh, shank, and heel of each lower limb (Fig. 1B) (O’Neill et al., 2015). Subjects were instructed to walk at a comfortable pace for 200 m (five back-and-forth laps, each 20 m length) in the lab to assess their preferred gait speed. Gait speed was measured using two photoelectric sensors placed 6 m apart in the middle of the walkway. The 6 m speed was recorded during the last four laps of the 200 m walk and averaged across the four trials to calculate the preferred gait speed. The measurement of preferred speed and all subsequent experimental trials were performed barefoot by each subject, to match the chimpanzee conditions.

**Fig. 1.**
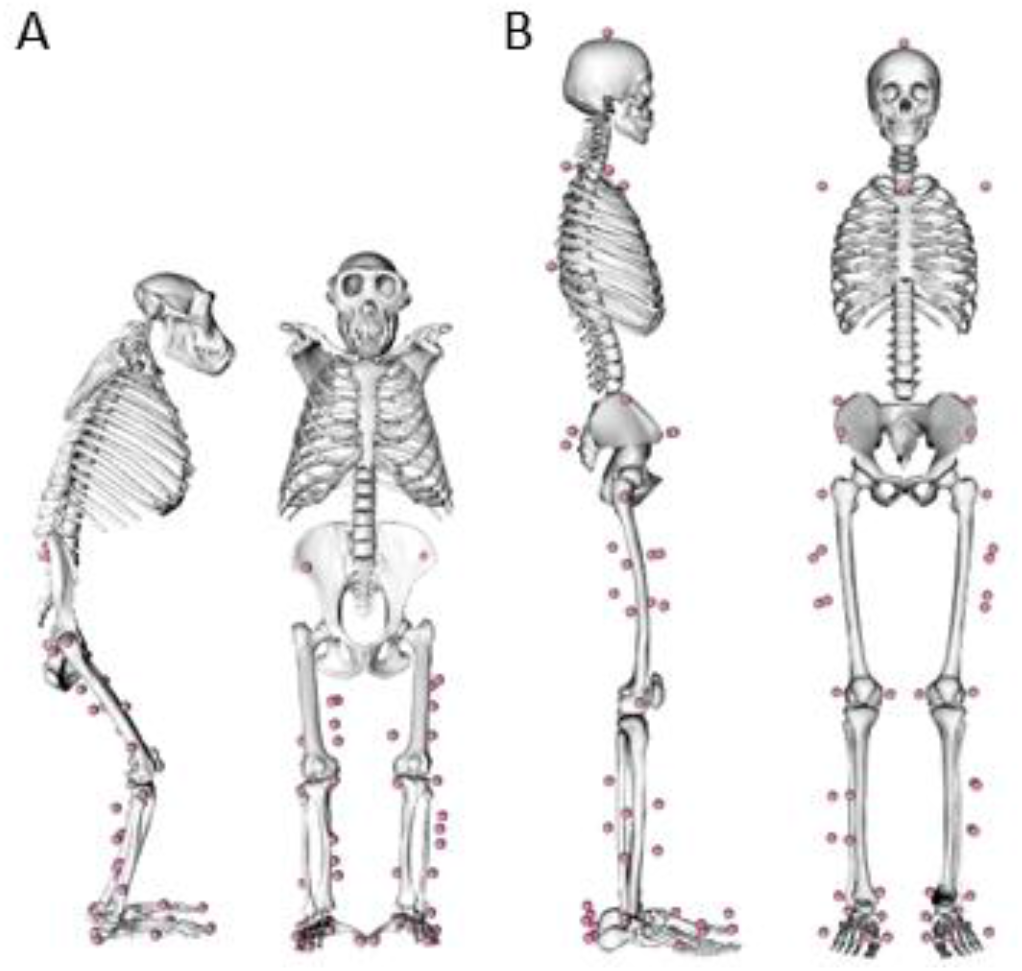
Musculoskeletal Morphology and Marker Placement. Representations of the skeletal morphology and marker positions (A) for the chimpanzee model, along with the skeletal morphology and marker positions (B) for the human model in OpenSim. The anatomical markers and segment marker clusters are depicted with pink dots.

Data were collected from the human subjects walking under four different postural conditions to facilitate comparisons with existing chimpanzee gait (O’Neill et al., 2015; O’Neill et al., in review). The four human postural conditions were designated: (1) normal (Norm), (2) crouched-limb (CL), (3) crouched-limb, flexed-trunk (CLT), and finally (4) crouched-limb, flexed-trunk, pelvis list (CLTP). The CLTP condition involved providing instructions specific to replicating the frontal plane pelvis motion observed in chimpanzees, which is an out-of-phase pattern compared with typical normal human gait (O’Neill et al., 2015). Each condition was performed at two speeds: the preferred gait speed and at a speed that matched the average, absolute walking speed for the previously collected chimpanzee data (chimpanzee-matched speed = 1.09 m/s). The order of the postural conditions was performed in sequence due to the additive nature of the instructions; however, the order of the two speeds was balanced among subjects within each condition (five did the chimpanzee-matched speed first for each condition).

Before beginning the walking trials, a static calibration trial was collected for the purpose of scaling a generic human musculoskeletal model to the size of each subject. For the Norm condition, subjects were instructed to “walk normally” at the given speeds. Next, the CL condition was meant to replicate, as best as possible, the designs from previous two-dimensional crouched posture studies that focused on sagittal plane variables (Carey and Crompton, 2005; Foster et al., 2013). Prior to the CL condition, the shoulder height of the subject was measured while standing crouched with the hip flexed 50° and the knee flexed 30° using a large goniometer. A taut rope was stretched along the walkway at this measured shoulder height. Subjects were instructed to “walk while bending at the hips and knees to match the target rope height” at the given speed. Next, the CLT condition was designed to target the difference in trunk angle between humans and chimpanzees during bipedal gait, and was based on the average bipedal chimpanzee trunk angle (Pontzer et al., 2014). The height of the rope was reset based on the shoulder height of the subjects while standing with the trunk flexed forward 30° from the vertical, the hip flexed 50°, and the knee flexed 30°. Subjects were instructed to “walk while bending at the trunk, hips, and knees to match the target rope height” at the given speed. For the final condition (CLTP), the height of the rope remained the same as in CLT and each subject was instructed to “walk like you did in the previous condition, but now focus on pitching your trunk and pelvis over the supporting limb during the swing phase.” Subjects also viewed a short, animated video of a chimpanzee walking bipedally to help them understand the desired pelvis list pattern. This condition was specifically designed to address the out-of-phase pelvis list motion in chimpanzees relative to normal human gait (O’Neill et al., 2015).

Each of the eight conditions (four postures times two speeds) were performed by the subjects as they walked overground across a walkway with three embedded force platforms (AMTI, Watertown MA, USA) while GRFs were recorded at 1200 Hz. Kinematic data were collected simultaneously at 240 Hz using an eleven-camera motion capture system (Qualisys, Gothenburg, Sweden). Both the 3-D raw marker trajectories and GRF data were smoothed using a fourth order zero-lag Butterworth low-pass filter with a cut-off frequency of 6 Hz.

Before each of the conditions, subjects were instructed to practice each walking task for at least three bouts and had the opportunity to ask questions about the instructions. The practice bouts were deemed to be completed once subjects were readily able to reproduce the specified posture and speed. Following the practice bouts, three acceptable trials were recorded for each of the eight conditions. A trial was considered acceptable if: (i) the gait speed was within 3% of the target speed, (ii) the feet cleanly struck each of the three force plates in the correct sequence, and (iii) the subject maintained the target posture throughout the trial. Maintenance of the target posture was assessed visually, with the help of the rope stretched along the walkway. Subjects were given the opportunity to rest between trials to minimize the chance of becoming fatigued during the data collection. Subjects self-reported that they did not become fatigued by the end of the data collection.

A 3-D model (Lai et al., 2017) of the human musculoskeletal system was used to determine the kinematic patterns using OpenSim software (Delp et al., 2007; Seth et al., 2018). The human model had 21 mechanical degrees-of-freedom (DOF), with a six DOF pelvis segment that articulates with a rigid segment representing the head, arms, and trunk via a ball-and-socket joint. Pelvis angles were calculated in the global reference frame relative to a neutral position for the human and chimpanzee gait data, and were calculated with a rotation-list-tilt rotation sequence (Baker, 2001). Each lower limb articulated to the pelvis via a ball-and-socket hip joint, the knee was represented as a modified hinge joint that included the translation of the tibia relative to the femur (Yamaguchi and Zajac, 1989), while the ankle and metatarsophalangeal joints were modeled as hinge joints. Lastly, to compare human trunk kinematics between postural conditions and confirm that the subjects were performing the tasks as instructed, trunk segment angles were calculated relative to the horizontal global reference frame.

For each of the ten subjects, a generic model was scaled to best match the individual anthropometrics of the subject based on the static, standing calibration trial. The scaled model was then used to calculate the generalized coordinates for each trial using the inverse kinematics algorithm in OpenSim (Lu and O’Connor, 1999). The pelvis rotation, pelvis list, pelvis tilt, hip flexion, hip adduction, hip rotation, knee flexion, and ankle flexion angles from the inverse kinematics analysis, and the vertical, anterior-posterior, and medial-lateral GRFs were compared with corresponding chimpanzee bipedal gait data.

### Chimpanzee Protocol

The chimpanzee kinematic and GRF data were drawn from previously published studies (O’Neill et al., 2015; O’Neill et al., in review). Therein, three male common chimpanzees (*Pan troglodytes*; age: 5.5 ± 0.2 years; mass: 26.5 ± 6.7 kg; hind limb length: 0.39 ± 0.02 m) walked overground while 3-D kinematic data were recorded at 150 Hz and GRF data were recorded at 1500 Hz. The 3-D marker trajectories were filtered using a fourth order zero-lag Butterworth low-pass filter, with cut-off frequencies set within the range of 4-6 Hz based on visual inspection of the data and the GRFs were low-pass filtered at 60 Hz. Each chimpanzee was trained to walk bipedally using a food reward and positive reinforcement for at least six months before data collection began. Four trials were collected per chimpanzee, while they walked at a self-selected speed following an animal trainer offering a food reward. Following data collection, a generic chimpanzee musculoskeletal model (O’Neill et al., 2013) was scaled to each individual chimpanzee using a short series of video frames obtained during the double support period. The chimpanzee musculoskeletal model had 16 mechanical DOF representing the hind limbs and pelvis segment (Fig. 1A). The mechanical DOF in the chimpanzee model were consistent with the human musculoskeletal model for the pelvis, hips, knees, and ankles. To compare with the human data, the corresponding chimpanzee angles and GRFs were used for the statistical analyses.

### Output Variables and Statistical Analyses

For each of the trials for each of the subjects, the following variables were calculated: stride length was calculated as the anterior-posterior pelvis displacement during the stride, stride time was the total time that it took to complete the stride, and gait speed was the stride length divided by stride time. Step width for each trial was calculated as the medial-lateral distance between the average centers of pressure under each foot during stance phase (Donelan et al., 2001). The minimum and maximum joint angles were calculated from the average subject data for each condition, and the ROM was calculated as the difference between the minimum and maximum joint angle.

The eight kinematic variables and three GRF variables described in the previous sections were compared for each of the four different human posture conditions to the chimpanzee bipedal gait mechanics (for both speeds: Norm vs Chimpanzee, CL vs. Chimpanzee, CLT vs. Chimpanzee, and CLTP vs. Chimpanzee). The kinematic data for each trial were time normalized to the full stride, and GRF data for each trial were time normalized to the stance phase. The time normalized data were averaged across trials for each condition for each subject, then group averages were calculated for each condition. The timing of GRF data were normalized to the stance phase, rather than the full stride, so that the comparisons would not be affected by the swing phase when all forces are zero. The magnitude of GRF data was normalized to body weight. Similarities in pattern and differences in magnitude for the kinematic and GRF variables were evaluated using zero-lag cross-correlations (*r*) and root-mean-square deviations (RMSD) based on the group average data.

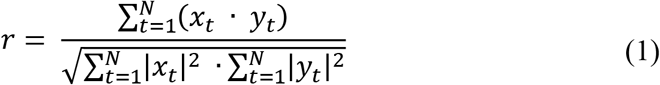

and

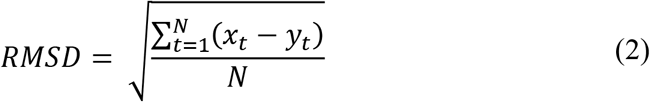

where the segment, joint, or GRF values at time *t* is given as *x_t_* (humans) and *y_t_* (chimpanzees) with *N* data points (*N* = 101). Values for *r* could vary from −1 to 1, and a greater positive value of *r* (closer to 1) would indicate that two variables were more similar to each other in pattern throughout the gait cycle. The minimum value for RMSD is 0, which would indicate that the variables have the same value throughout the entire gait cycle, while greater RMSD values indicate greater differences in magnitude between the two species. Four unique outcome measures were computed for each condition to assess the similarity between human and chimpanzee gait mechanics: i) the average of *r* across the eight kinematic variables, ii) the average of RMSD across the eight kinematic variables, iii) the average of *r* for the three orthogonal components of the GRFs, and iv) the average of RMSD for the three orthogonal components of the GRFs.

## Results

The average preferred gait speed for the ten human subjects was 1.30 ± 0.15 m s^−1^. During the chimpanzee-matched-speed trials, the human subjects were able to closely match (1.10 ± 0.02 m s^−1^) the average chimpanzee bipedal gait speed (O’Neill et al. 2015; 1.09 m s^−1^; Table 1). There were only subtle differences in kinematics and GRFs between the two human speeds for each posture condition, therefore only the results comparing the chimpanzee gait data to human gait at the chimpanzee-matched speed are reported in the following sections (see supplementary material for corresponding results for human preferred speed to chimpanzee gait). In the spatiotemporal domain, the chimpanzees took shorter and quicker strides than the humans across all posture conditions (Table 1 and Table S1). The stance time, stride length, and stride frequency were similar across the human posture conditions. Chimpanzees walked with a greater step width (0.26 ± 0.04 m) than any of the human conditions, both in absolute terms and relative to hind limb length. However, the humans had wider steps in the crouched posture conditions than the normal condition, with the greatest difference being CLTP (0.21 ± 0.07 m) versus Norm (0.14 ± 0.03 m).

**Table 1:** Spatiotemporal results for the four human conditions (chimpanzee-matched speed) and the chimpanzee bipedal gait (mean ± standard deviation).

|  | velocity (m/s) | stance time (s) | swing time (s) | stride length (m) | stride freq. (Hz) | step width (m) | duty factor |
| --- | --- | --- | --- | --- | --- | --- | --- |
| Chimp. | $1.11 \pm 0.06$ | $0.46 \pm 0.07$ | $0.28 \pm 0.01$ | $0.81 \pm 0.04$ | $1.43 \pm 0.23$ | $0.26 \pm 0.04$ | $0.62 \pm 0.03$ |
| Normal | $1.11 \pm 0.03$ | $0.73 \pm 0.04$ | $0.40 \pm 0.02$ | $1.24 \pm 0.06$ | $0.89 \pm 0.04$ | $0.14 \pm 0.03$ | $0.65 \pm 0.01$ |
| CL | $1.10 \pm 0.02$ | $0.75 \pm 0.04$ | $0.37 \pm 0.03$ | $1.23 \pm 0.05$ | $0.91 \pm 0.06$ | $0.17 \pm 0.04$ | $0.67 \pm 0.01$ |
| CLT | $1.10 \pm 0.02$ | $0.75 \pm 0.04$ | $0.37 \pm 0.03$ | $1.23 \pm 0.06$ | $0.90 \pm 0.05$ | $0.17 \pm 0.05$ | $0.67 \pm 0.01$ |
| CLTP | $1.10 \pm 0.02$ | $0.77 \pm 0.05$ | $0.39 \pm 0.04$ | $1.27 \pm 0.09$ | $0.88 \pm 0.06$ | $0.21 \pm 0.07$ | $0.66 \pm 0.02$ |

### Pelvis kinematics

Both the human and chimpanzee subjects internally rotated their pelvis during the first half of the gait cycle, followed by external pelvis rotation during the second half of the gait cycle (Fig. 2A and 2B). However, the chimpanzees had greater pelvis rotation ROM (46 ± 12°) over the gait cycle than for any of the human conditions (Table 2). The pelvis rotation ROM was greatest in the CLTP condition than the other human conditions (24 ± 9°). When assessing the similarity of the human conditions to the chimpanzee gait using the cross-correlations and RMSD, the CLTP human posture conditions produced the most similar pelvis rotation motion in pattern (*r* = 0.61) and magnitude (RMSD = 10.8°) compared to chimpanzee gait (Table 3).

**Fig. 2.**
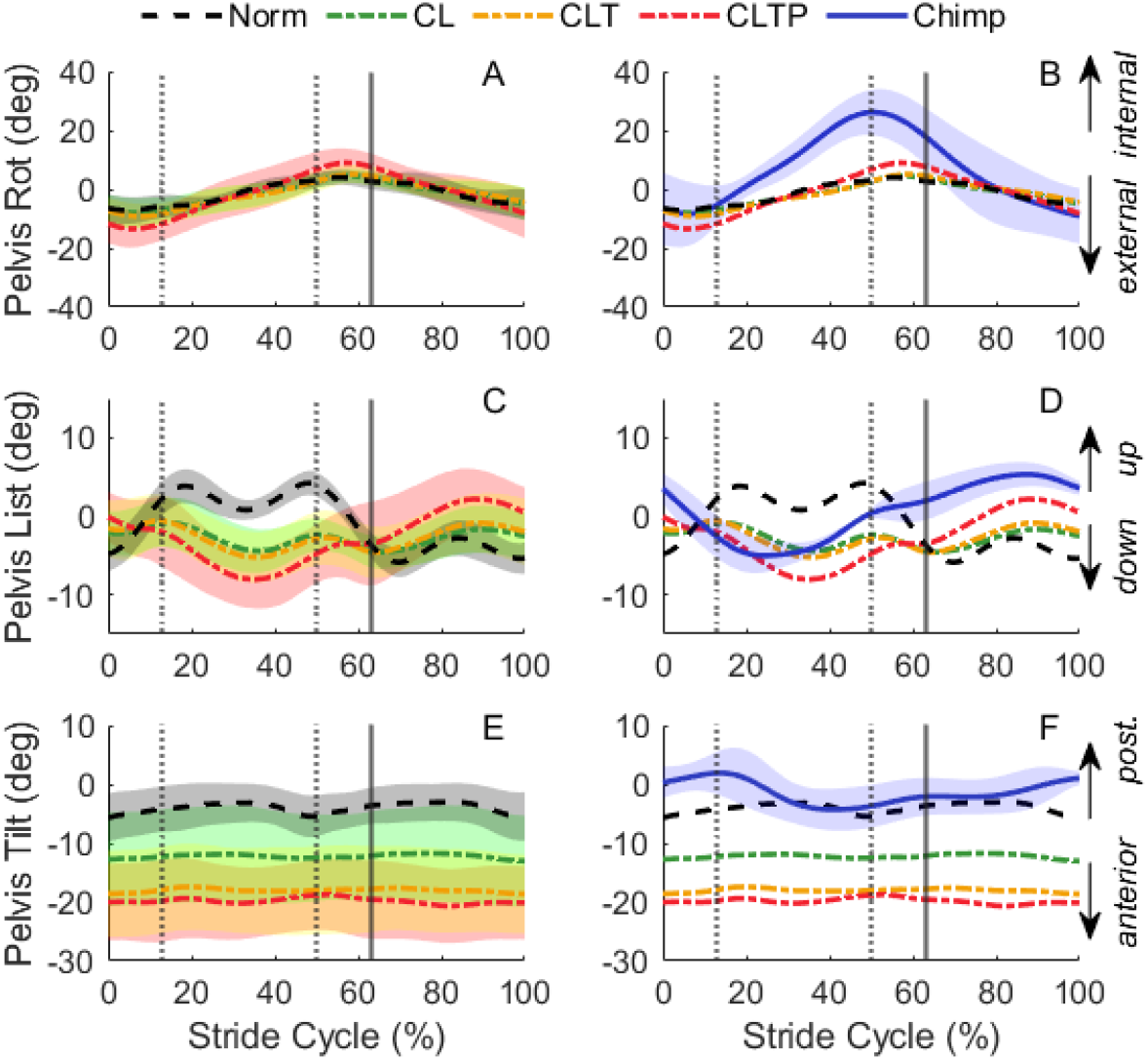
Pelvis Kinematics. The left column contains the pelvis rotation (A) list (C) and tilt (E) kinematics for each of the four human conditions at the chimpanzee-matched speed (Norm, CL, CLT, CLTP), with the shaded regions representing 1 standard deviation of the mean. The right column shows the means for the pelvis rotation (B), list (D), and tilt (F) for each of the four human conditions along with the mean and standard deviation for the chimpanzee data (Chimp). The vertical lines depict left toe-off, left foot strike, and right toe-off respectively. Table S1 provides exact gait events across conditions.

**Table 2:** Segment and joint angle minimums (Min), maximums (Max), and range of motion (ROM) for each of the conditions (units in degrees; mean ± standard deviation).

|  |  | Pelvis Rot. | Pelvis List | Pelvis Tilt | Hip Flex. | Hip Add. | Hip Rot. | Knee | Ankle |
| --- | --- | --- | --- | --- | --- | --- | --- | --- | --- |
| Normal | Min | -5 ± 3 | -5 ± 2 | -7 ± 4 | -10 ± 6 | -11 ± 3 | -13 ± 4 | 5 ± 2 | -14 ± 5 |
| | Max | 8 ± 4 | 7 ± 2 | -2 ± 3 | 31 ± 4 | 8 ± 2 | -2 ± 4 | 64 ± 4 | 16 ± 4 |
| | ROM | 12 ± 5 | 12 ± 3 | 4 ± 1 | 42 ± 2 | 19 ± 4 | 11 ± 3 | 59 ± 2 | 30 ± 4 |
| CL | Min | -5 ± 2 | 0 ± 3 | -14 ± 8 | 18 ± 18 | -10 ± 5 | -9 ± 3 | 32 ± 6 | 0 ± 5 |
| | Max | 9 ± 4 | 6 ± 2 | -11 ± 8 | 58 ± 12 | -1 ± 4 | 2 ± 3 | 86 ± 7 | 34 ± 5 |
| | ROM | 14 ± 4 | 6 ± 2 | 3 ± 1 | 40 ± 8 | 9 ± 4 | 11 ± 3 | 55 ± 7 | 35 ± 5 |
| CLT | Min | -6 ± 3 | 0 ± 3 | -20 ± 7 | 30 ± 11 | -10 ± 5 | -10 ± 4 | 32 ± 2 | -2 ± 5 |
| | Max | 10 ± 4 | 6 ± 4 | -16 ± 7 | 66 ± 7 | -1 ± 5 | 4 ± 4 | 90 ± 7 | 35 ± 6 |
| | ROM | 15 ± 6 | 6 ± 3 | 3 ± 1 | 36 ± 5 | 9 ± 3 | 13 ± 4 | 59 ± 6 | 37 ± 6 |
| CLTP | Min | -10 ± 5 | -3 ± 3 | -22 ± 6 | 29 ± 10 | -15 ± 6 | -13 ± 5 | 34 ± 5 | -3 ± 7 |
| | Max | 14 ± 6 | 9 ± 5 | -18 ± 6 | 68 ± 8 | -5 ± 5 | 2 ± 5 | 90 ± 8 | 35 ± 8 |
| | ROM | 23 ± 9 | 12 ± 4 | 4 ± 2 | 40 ± 7 | 9 ± 3 | 16 ± 4 | 56 ± 8 | 39 ± 9 |
| Chimp. | Min | -12 ± 7 | -6 ± 1 | -5 ± 4 | 25 ± 9 | -30 ± 3 | -35 ± 3 | 14 ± 3 | -19 ± 8 |
| | Max | 29 ± 12 | 6 ± 1 | 3 ± 3 | 52 ± 6 | -14 ± 7 | 4 ± 4 | 92 ± 2 | 19 ± 4 |
| | ROM | 41 ± 13 | 12 ± 2 | 8 ± 1 | 27 ± 4 | 16 ± 4 | 39 ± 2 | 78 ± 1 | 38 ± 5 |

**Table 3:**
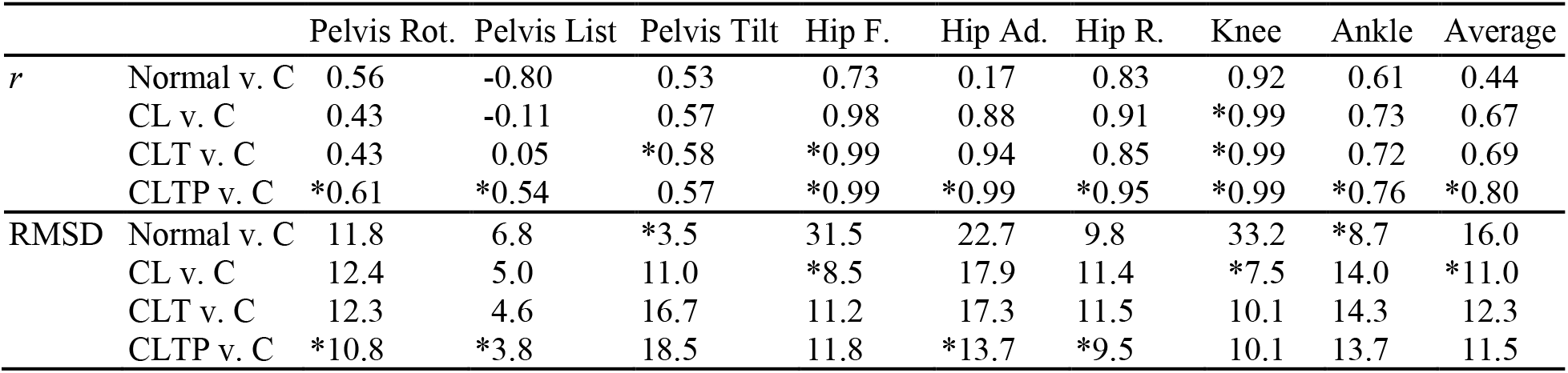
Cross-correlation coefficients (*r*) and RMSD (units in degrees) for kinematics for humans walking with chimpanzee-matched speed (1.11 ± 0.02 m/s) compared with chimpanzee gait (C); asterisks indicate human condition that was most similar to chimpanzee gait.

**Table 4:**
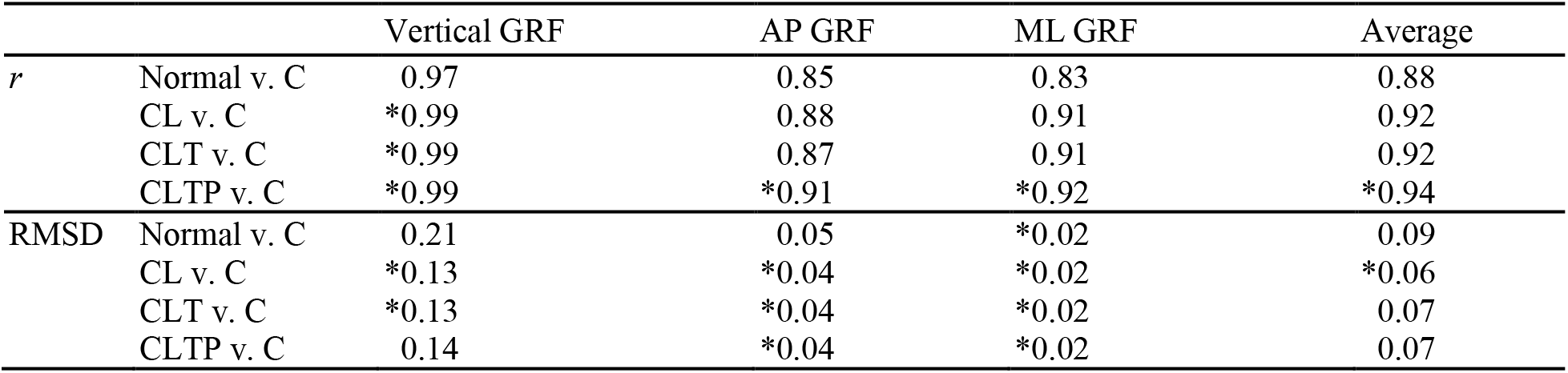
Cross-correlation coefficients (*r*) and RMSD (units in BW) for GRFs for humans walking with chimpanzee-matched speed (1.11 ± 0.02 m/s) compared with chimpanzee gait (C); asterisks indicate human condition that was most similar to chimpanzee gait.

|  |  | Vertical GRF | AP GRF | ML GRF | Average |
| --- | --- | --- | --- | --- | --- |
| $r$ | Normal v. C | 0.97 | 0.85 | 0.83 | 0.88 |
|  | CL v. C | *0.99 | 0.88 | 0.91 | 0.92 |
|  | CLT v. C | *0.99 | 0.87 | 0.91 | 0.92 |
|  | CLTP v. C | *0.99 | *0.91 | *0.92 | *0.94 |
| RMSD | Normal v. C | 0.21 | 0.05 | *0.02 | 0.09 |
|  | CL v. C | *0.13 | *0.04 | *0.02 | *0.06 |
|  | CLT v. C | *0.13 | *0.04 | *0.02 | 0.07 |
|  | CLTP v. C | 0.14 | *0.04 | *0.02 | 0.07 |

In the frontal plane during normal gait, the human pelvis listed downward towards the swing side limb, whereas the chimpanzees elevated their pelvis on the swing side (Fig. 2C and 2D). The out-of-phase pattern of motion between the normal human condition and chimpanzees is reflected in the −0.80 *r* value (Table 3). The pelvis list ROM was smaller in the CL and CLT posture conditions than the normal condition (Table 2), and this relatively straight-line pelvis list trajectory resulted in *r* values that were close to zero for these conditions (CL: *r* = −0.11; CLT: *r* = 0.05). For the CLTP condition, the human pelvis list pattern was in-phase with the chimpanzee data (*r* = 0.54) and had the most chimpanzee-like magnitude (RMSD = 3.8°; Table 3).

Humans tilted their pelvis slightly forward for the normal posture condition, between −2° and −7°, throughout the gait cycle (Fig. 2E and 2F; Table 2). Chimpanzees also walked with a relatively vertical orientation of the pelvis, which resulted in a close match of the RMSD between the normal human condition and the chimpanzees (Norm: RMSD = 3.5°; Table 3). The different human posture conditions had little effect on the patterns of motion as reflected by the *r* values (*r* = [0.53, 0.58]; Table 3). However, during the crouched posture conditions humans tilted their pelvis forward to a greater extent (CL: peak = −14°; CLTP: peak = −22°; Table 2) than for the normal condition. This resulted in greater RMSD values for the crouched human posture conditions than the normal condition (CLTP: RMSD = 18.5°; Table 3). In summary, the pelvis rotation and pelvis list motions were most chimpanzee-like in the CLTP condition, however the pelvis tilt kinematics were less chimpanzee-like in the crouched posture conditions compared to the normal human condition.

### Lower/hind limb kinematics

In the normal human posture condition, humans began in hip flexion, gradually extended the hip throughout the stance phase to hyperextension, then gradually flexed the hip again during the swing phase. During the human crouched posture conditions, the hip was more flexed throughout the stride compared to the normal human condition (Table 2; Fig. 3A and 3B). Hip flexion was more similar in pattern (CLTP: *r* = 0.99) and magnitude (CL: RMSD = 8.5°) to the chimpanzees in the human crouched posture conditions than during the normal human condition (Table 3). There were only small differences in hip flexion motion across the different human crouched posture conditions.

**Fig. 3.**
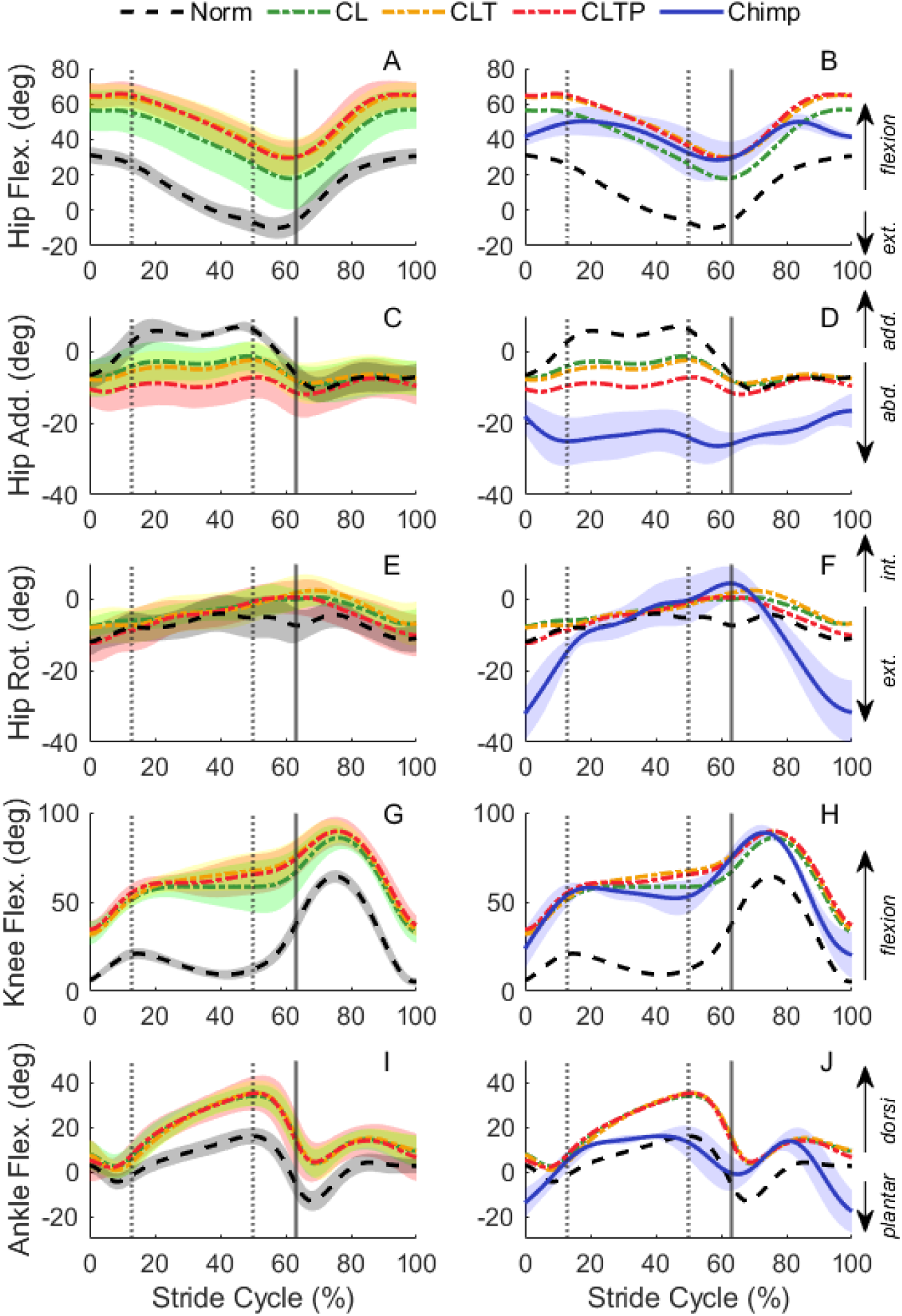
Hind limb kinematics. The left column shows the hip flexion (A) hip adduction (C), hip rotation (E), knee flexion (G) and ankle flexion (I) kinematics for each of the four human conditions at the chimpanzee-matched speed (Norm, CL, CLT, CLTP), with the shaded regions representing 1 standard deviation of the mean. The right column shows the means for the hip flexion (B) hip adduction (D), hip rotation (F), knee flexion (H) and ankle flexion (J) kinematics for each of the four human conditions along with the mean and standard deviation for the chimpanzee data (Chimp).

In the frontal plane, humans began the gait cycle in an abducted hip position, adduct during the early part of stance phase, then abduct during the late stance phase (Fig. 3C and 3D). In contrast, chimpanzees maintained their hip in an abducted position throughout the stance and swing phases with a magnitude that exceeded the peak human hip abduction angle. Across the human crouched posture conditions, humans had greater hip abduction than the normal human condition, although it did not reach the same magnitude of hip abduction as the chimpanzees (peak abduction angle: chimpanzees = −31 ± 4°; human CLTP = −15 ± 6°). The CLTP condition produced the most similar hip abduction pattern (*r* = 0.99) and magnitude (RMSD = 13.7°) to the chimpanzees (Table 3).

Both humans and chimpanzees internally rotated their hip during the stance phase and externally rotated their hip during swing (Fig. 3E and 3F). However, as was seen for the pelvis rotation, the chimpanzees had greater ROM (42 ± 3°) than any of the human conditions (greatest ROM for CLTP = 17 ± 4°). There were only subtle differences in hip rotation across the human posture conditions, with the CLTP condition producing the most chimpanzee-like hip rotation patterns (*r* = 0.95) and magnitude (RMSD = 9.5°; Table 3).

In the normal human condition, the humans had a knee angle close to full extension throughout most of the stance phase, then flexed the knee rapidly in preparation for swing phase. Chimpanzees had a more flexed knee angle throughout the gait cycle than the normal human condition (Fig. 3G and 3H). The knee angle pattern was more similar to chimpanzees in each of the human crouched posture conditions (*r* = 0.99; Table 3) than the normal human condition. The magnitude of knee angle in the crouched posture conditions (CL: RMSD = 7.5°) was also more similar to the magnitude of the chimpanzee knee angle than the normal human condition (RMSD = 33.2°).

In the normal human condition, humans rapidly plantar flexed the ankle early in the stance phase, gradually dorsiflexed the ankle throughout midstance, followed by a rapid plantar flexion motion during push-off phase (Fig. 3I and 3J). The ankle angle patterns were more chimpanzee-like in the human crouched posture conditions than the normal condition (Norm: *r* = 0.61; CLTP: *r* = 0.76). However, the ankle was more dorsiflexed throughout stance phase in the crouched posture conditions than the normal condition, resulting in a greater RMSD for the crouched posture conditions (Norm: RMSD = 8.7°; CLTP: RMSD = 13.7°).

Overall, the kinematics patterns (measured by *r*) were more chimpanzee-like in the human crouched posture conditions than the normal human condition. However, the magnitudes of the differences in kinematics (measured by RMSD) had different responses across variables, with the normal human condition resulting in more chimpanzee-like pelvis tilt and ankle angle magnitudes than the human crouched posture conditions.

### Trunk Kinematics

Human subjects maintained an upright trunk segment angle in normal gait (Fig. 4A), while in the CL condition, humans chose to lean their trunk forward ~25° from vertical. When given specific instructions to tilt forward at the trunk in the CLT and CLTP conditions, the human subjects leaned forward to ~35° on average. For each of the CL, CLT, and CLTP conditions, the forward flexion of the trunk was achieved through a combination of lumbar flexion (Fig. 4B) and pelvis tilt (Fig. 2E)

**Fig. 4.**
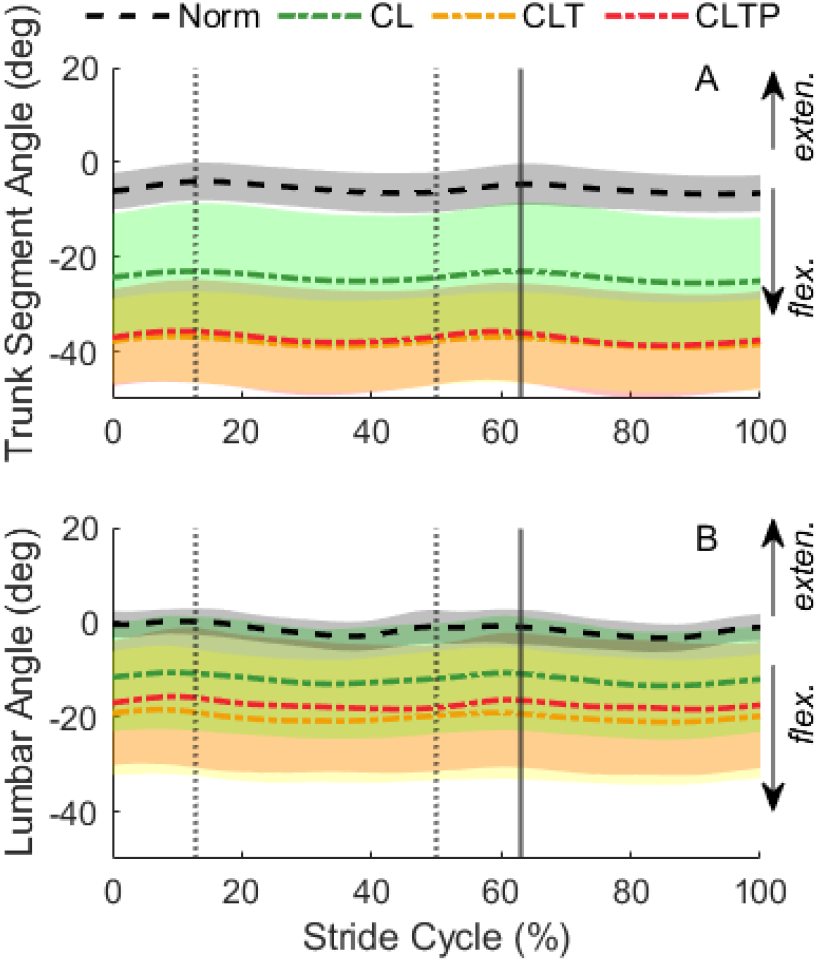
Lumbar Joint and Trunk Segment Angles. The trunk segment angles (A) and lumbar joint angles (B) for each of the four human conditions for the chimpanzee-matched speed. Trunk angles were calculated as the sum of pelvis tilt and lumbar extension.

### Ground Reaction Forces

The vertical GRF had a double-peaked pattern in the normal human condition (Fig. 5A and 5B). During the human crouched posture conditions, the vertical GRF still had a double-peaked shape but had a reduced second peak, and less of a trough between the peaks, compared to the normal human condition. The average chimpanzee vertical GRF pattern had only one distinct peak, which occurred in the first half of stance phase. According to the cross-correlation coefficient, the pattern of the vertical GRFs were similar in both the normal human condition (*r* = 0.97), and each of the crouched human postures (*r* = 0.99). The magnitude difference for the vertical GRFs were greater in the normal condition than the crouched posture conditions (Norm: RMSD = 0.21; CL: RMSD = 0.13), with only subtle differences in the magnitude of the vertical GRF across the different human crouched posture conditions.

**Fig. 5.**
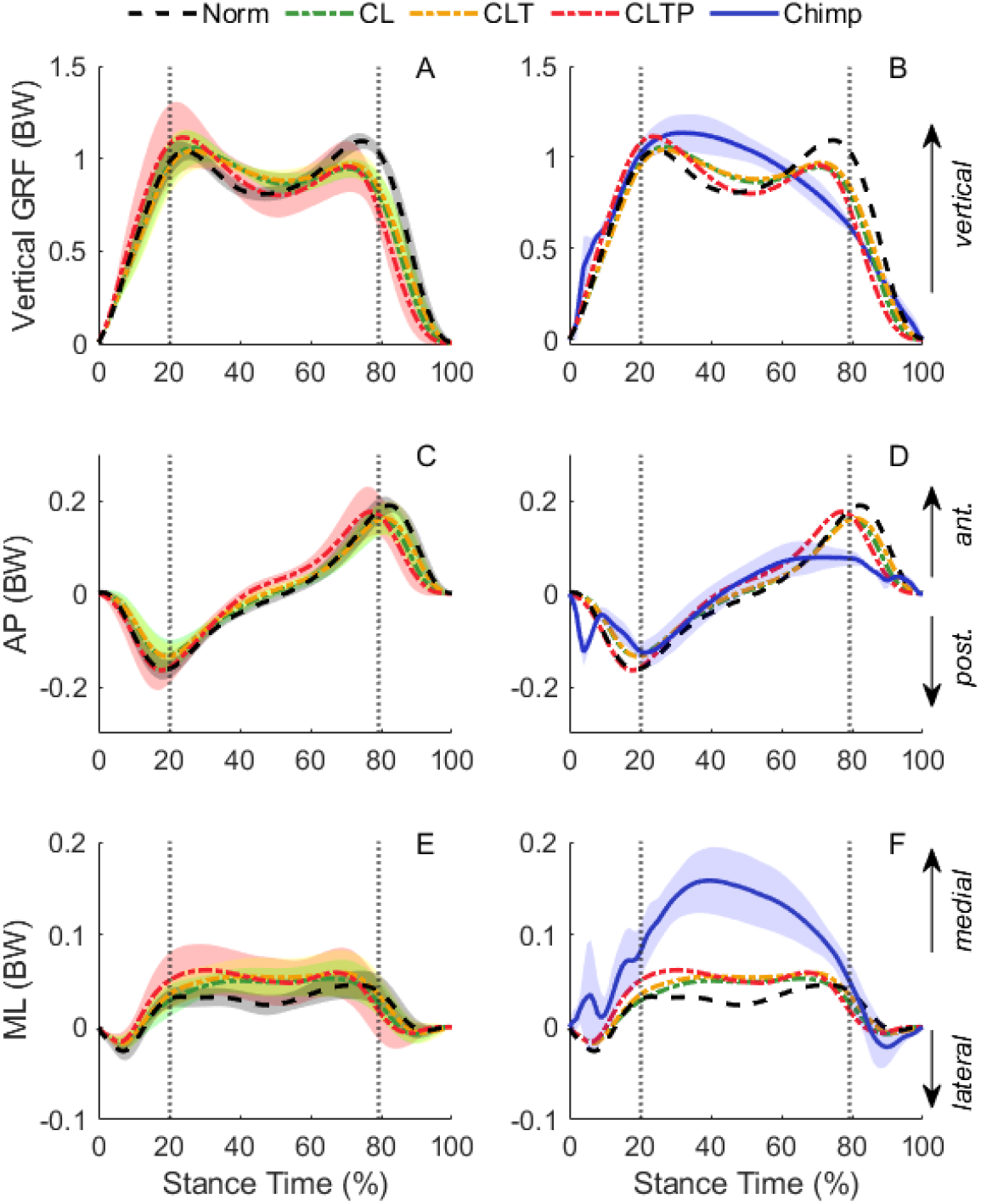
Ground reaction forces (GRFs). The left column shows the GRFs for the vertical (A) anterior-posterior (C), and medial-lateral (E) directions for each of the four human conditions at the chimpanzee-matched speed (Norm, CL, CLT, CLTP), with the shaded regions representing 1 standard deviation of the mean. The right column shows the means for the GRFs for the vertical (B) anterior-posterior (D), and medial-lateral (F) directions for each of the four human conditions along with the mean and standard deviation for the chimpanzee data (Chimp).

The gait of both humans and chimpanzees had a negative, posteriorly directed GRF during the first half of stance phase, then a positive, anteriorly directed GRF for the second half of stance phase (Fig. 5C and 5D). The push-off force during the second half of stance was different between humans and chimpanzees, as the peak positive GRF for the human conditions were greater than the chimpanzees. When compared with the normal human condition, the human crouched posture conditions produced only slightly more chimpanzee-like AP GRF patterns (Norm: *r* = 0.85; CLTP: *r* = 0.91) and magnitudes (Norm: RMSD = 0.05; CLTP: RMSD = 0.04).

In the normal human condition, the ML GRF had a negative, laterally directed peak during early stance and then a positive, medially directed GRF throughout midstance (Fig. 5E and 5F). The human crouched posture conditions produced greater medial GRFs throughout midstance than the normal human conditions, but the peak magnitude did not approach the magnitude for the chimpanzees. When compared with each of the human conditions, chimpanzees produced greater medial GRFs throughout the middle of stance phase. The ML GRF was more chimpanzee-like in the human crouched posture conditions than the normal human condition (Norm: *r =*0.83; CLTP: *r* = 0.92). The RMSD magnitudes did not vary across any of the human posture conditions (RMSD = 0.02). Overall, in all three planes of GRF, the GRFs were more chimpanzee-like in the human crouched postures than the normal human condition, but the human crouched postures did not produce a mono-phasic vertical GRF as was observed in chimpanzees.

### Average r and RMSD for Kinematics and GRFs

Each of the human crouched posture conditions produced more chimpanzee-like gait mechanics than the normal human condition according to each of the four metrics used to evaluate similarity (average *r* and RMSD for the kinematics and GRFs, respectively). Thus, the instructions provided to the subjects were generally effective at producing more chimpanzee-like gait mechanics than normal human gait. When evaluating how closely the three different crouched posture conditions replicated chimpanzee gait, there were fewer generalizations that could be drawn. For the kinematic patterns, evaluated by the average *r*, the kinematic patterns in the CLTP condition were more chimpanzee-like than the other three conditions. The hip flexion, hip adduction, and hip rotation had *r* values greater than 0.95 in the CLTP condition, which indicates a high similarity in pattern with the chimpanzee data for the hip angles. Compared with the CL condition, the CLT condition with the forward trunk flexion instruction, did not further increase *r* values. For the other average metrics (kinematic RMSD, GRF *r* and RMSD), there was no single condition that most closely matched the chimpanzee data across all metrics. Additionally, for some variables such as pelvis rotation and ankle flexion, the crouched posture conditions did not result in a close match to the chimpanzee data in *r* or RMSD. Thus, while the crouched posture instructions produced a more chimpanzee-like gait than normal human gait overall, the three crouched posture conditions did not result in additive changes to gait mechanics that progressively better matched the bipedal chimpanzee mechanics, as hypothesized.

## Discussion

We compared the 3-D gait mechanics of normal walking and crouched posture walking in humans to bipedal walking in chimpanzees to understand the degree to which human crouched posture walking is similar to that of bipedal gait in chimpanzees. Our first hypothesis was that the normal human condition would be least like the chimpanzee gait mechanics compared with any of the crouched posture conditions. Our data support this hypothesis as the normal human walking condition had the smallest average *r* and greatest average RMSD values for the kinematics and GRFs compared with each of the other human crouched postures. While each of the human crouched posture conditions produced more chimpanzee-like gait patterns than normal human walking, there was not strong support for the hypothesis that the human CLT condition would be more similar to chimpanzees than the human CL condition, as the gait mechanics between these conditions were similar. Our final hypothesis was that the joint kinematics and GRFs from the human CLTP condition would be most similar to the chimpanzee gait mechanics. The results for this hypothesis were more nuanced. The human CLTP condition did result in the most chimpanzee-like kinematic patterns (as measured by *r*). However, all three crouched posture conditions had similar patterns (*r*) for the GRFs, and similar magnitude differences (RMSD) for both kinematics and GRFs. The greater average *r* value for the kinematic variables in the CLTP condition than the CL or CLT conditions was mostly driven by a more chimpanzee-like pelvis list pattern, corresponding with the instructions to modify the pelvis list motion.

### Pelvis Kinematics

Across all human posture conditions, one of the persistent kinematic differences between humans and chimpanzees was the pelvis transverse plane rotation ROM (Fig. 2B; Table 2). On average, chimpanzees had 41° of pelvis rotation ROM, while humans had a pelvis rotation ROM between 12° and 23° across the four conditions. One potential explanation for this consistent interspecific difference is that chimpanzees have shorter hind limbs than humans in both an absolute and relative sense, so chimpanzees might use greater pelvis rotation throughout the gait cycle to increase their overall stride length (O’Neill et al., 2015; Whitcome et al., 2017). However, humans did not increase their pelvis rotation ROM when walking with the CL or CLT postures, even though these postures shorten their effective limb length (i.e., distance from the hip to the ground). The pelvis rotation ROM was moderately greater in the CLTP posture relative to the other crouched limb postures (i.e., CLPT: 23±9° ROM; CLT: 15±6° ROM), but that was still half of the measured pelvis rotation ROM for bipedal chimpanzees. Also, humans were free to modulate their stride length during each of the posture conditions, yet they used a consistent stride length across all four posture conditions (Table 1). Therefore, our data suggest that greater pelvis rotation ROM in chimpanzees than humans may not be a direct result of walking with shorter limbs or a crouched posture. Instead, the differences in pelvis rotation may reflect differences in the musculoskeletal design of the pelvis between humans and chimpanzees, such as orientation of the ilia and the corresponding effects on muscle function (e.g., Lovejoy, 2005; Lycett and von Cramon-Taubadel, 2013; Stern Jr. and Susman, 1983). Determination of individual muscle contributions to pelvis rotation in bipedal chimpanzees and humans would provide additional insight into this issue.

While the human crouched postures did not elicit chimpanzee-like pelvis transverse plane rotation, the pelvis list pattern was more chimpanzee-like across all human crouched postures compared with normal human gait, even though the CL and CLT instructions only targeted sagittal plane variables. The pelvis list patterns for normal human gait and bipedal chimpanzee gait are out-of-phase (Fig. 2D), but the pelvis list pattern in the human crouched posture conditions was in-phase with the chimpanzee pelvis list pattern. This indicates that an important determinate of raising the pelvis on the limb swing side is walking on a flexed lower limb, possibly to aid with foot-ground clearance (O’Neill et al., 2015, 2018). This also suggests that previous studies of human crouched posture walking likely produced a chimpanzee-like frontal plane pelvis motion, even if they did not measure it. However, the pelvis list motion was more chimpanzee-like in the CLPT condition than the CL or CLT conditions, which suggests that the pelvis list pattern of bipedal chimpanzees may be due to other factors beyond that of simply walking with a crouched posture. More human-based experiments could provide additional insight into the interaction (or lack thereof) between pelvis list and lower limb gait mechanics, perhaps by providing different instructions to humans to modify pelvis motion (e.g., Kikel et al., 2020), or by manipulating human anthropometrics (e.g., artificially increase foot length) which would affect foot-ground clearance during the swing phase of gait.

Humans walked with greater anterior pelvis tilt during all crouched posture conditions compared with normal walking (Fig. 2E). In the normal walking condition, the pelvis tilt exhibited by humans overlapped the chimpanzee data, with both generally falling within 5 of the upright, neural position. In contrast, the differences in pelvis tilt kinematics between humans and chimpanzees were greater in the crouched postures. The greater forward pelvis tilt used by humans in the crouched posture conditions, especially in the CL condition (tilted anteriorly 11-14°; Table 2), is more similar to bipedal macaques (tilted anteriorly 8-13°) than to bipedal chimpanzees (Ogihara et al., 2010; O’Neill et al., 2018).

### Lower/hind limb kinematics

The human sagittal plane hip and knee angles matched closely with the bipedal chimpanzee gait mechanics across each of the crouched posture conditions (Fig. 3B and 3H). Most of the previous crouched posture studies have focused on these two angles in particular, often referring to the gait pattern itself as ‘bent-hip, bent-knee” (Carey and Crompton, 2005; Crompton et al., 1998; Foster et al., 2013; Wang et al., 2003). Our data set is the first to directly compare joint kinematics of human crouched posture walking with bipedal chimpanzees, therefore these data provide important context for interpreting results of previous studies, especially our CL condition which replicates the experimental designs used in some prior research (Carey and Crompton, 2005; Foster et al., 2013). Nevertheless, the remaining differences observed in our data for both kinematics and GRFs illustrate that instructing humans to adopt a ‘bent-hip, bent-knee’ walking posture results in an overall gait pattern that is still quite different from bipedal chimpanzee walking in several important ways.

In normal human walking, the hip joint is adducted for most of the stance phase. In contrast, all three of the human crouched posture conditions resulted in an abducted hip position throughout the gait cycle. Additionally, humans had greater hip abduction during stance in the CLTP condition than the other crouched postures (Fig. 3C; Table 2), supporting previous results that swing side pelvis elevation (i.e. pelvis list) results in greater hip abduction (Kikel et al., 2020). However, the magnitude of the hip abduction angle across all crouched postures was still much less than that of chimpanzees (Fig. 3D). One potential explanation for the difference in the magnitude of the hip abduction angle is the presence of a frontal plane, valgus knee alignment in humans while chimpanzees lack this frontal plane angulation (Shefelbine et al., 2002; Tardieu and Trinkaus, 1994). A valgus knee enables humans to place their foot underneath the body center of mass throughout the stance phase, while maintaining an adducted hip position and with minimal upper body motion. The differences in knee alignment may allow the human subjects in this study to perform the crouched posture conditions with a lesser amount of hip abduction than the chimpanzees during bipedal gait. Likewise, *Australopithecus afarensis* had a valgus knee (Heiple and Lovejoy, 1971; Shefelbine et al., 2002), which suggests that regardless of the uncertainty in their posture (crouched vs. upright) they may have walked with less hip abduction than chimpanzees.

The differences in hip transverse plane rotation ROM between humans and chimpanzees persisted throughout all human crouched posture conditions. Humans averaged between 11 and 16° ROM across conditions, compared with 39° ROM for bipedal chimpanzees (Table 2). This persistent difference in hip rotation between humans and chimpanzees was similar to the persistent difference in pelvis transverse plane rotation for all human conditions compared with chimpanzees. However, one must be cautious in associating joint and segment motion when the femur is oriented ~45° relative to vertical in crouched limb walking, rather than being nearly vertical in upright walking. Specifically, pelvis transverse plane rotation, as well as pelvis list, will be directly influenced by both hip joint adduction and hip joint rotation in ways that are hard to decipher without conducting a 3-D kinematic analysis, as is reported here. For example, even with more chimpanzee-like pelvis list during human crouched posture walking, these conditions do not result in chimpanzee-like hip joint adduction or hip joint rotation in humans. This is functionally importantly, as hip joint orientation is what will impact the leverage and function of the hip muscles to provide balance and propulsion during walking (i.e., Stern, 1972; Stern and Susman, 1981). Future muscle-level analyses will be helpful in understanding how hip joint and pelvis motion are coordinated and actuated by muscle forces.

At the ankle, humans maintained a substantially more dorsiflexed ankle position throughout most of the stance phase in the crouched posture conditions than for either normal human walking or bipedal walking in chimpanzees (Fig. 3I and 3J). As with the pelvis tilt kinematics, the ankle joint represents another case where the human crouched posture conditions resulted in less chimpanzee-like kinematics than normal human walking, with likely implications for the function of ankle muscles. Due to the dorsiflexed ankle position at initial foot-ground contact in all conditions, humans contact the ground with their heel first (i.e., with a distinct heel strike) even when walking in the crouched postures. This early stance phase behavior is distinct from bipedal chimpanzees, who adopt a plantar flexed ankle angle at foot-strike and place the foot nearly flat on the ground at initial foot-ground contact (O’Neill et al., 2015). This implies that the previously noted difference in foot strike patterns between humans and bipedal chimpanzees (O’Neill et al., 2015; O’Neill et al., 2018; O’Neill et al., in review) are not simply a result of differences in lower limb flexion between species.

### Trunk Kinematics

There were only minor differences in the pelvis and lower limb kinematics between the CL and CLT conditions, demonstrating that an anteriorly flexed trunk does not substantially alter gait mechanics when humans walk with a crouched posture. While the trunk segment angles matched closely with our target trunk angle of −30° (Fig. 4A), humans achieved this segment angle through a combination of lumbar flexion (Fig. 4B) and pelvis tilt (Fig. 2E). The human subjects did this rather than maintaining a vertically oriented pelvis and flexing their vertebral column alone, as in bipedal chimpanzees (O’Neill et al., 2015). The incorporation of greater anterior pelvis tilt resulted in greater hip flexion angles throughout the gait cycle in the CLT condition than the CL condition. The lack of broad differences in kinematics between CL and CLT conditions agrees with previous research on the effect of a moderately flexed trunk angle on crouched gait mechanics (Grasso et al., 2000).

Other researchers have examined the isolated effect of trunk orientation on gait mechanics and found that an anteriorly flexed trunk angle of 30°, without any instructions to modify lower limb posture, resulted in greater knee flexion and ankle dorsiflexion angles throughout stance than normal walking (Aminiaghdam et al., 2017; Kluger et al., 2014; Saha et al., 2008). While the 30° trunk flexion gait resulted in a crouched limb posture in Aminiaghdam et al., more extreme changes in gait mechanics occurred at trunk flexion angles of 90°. Given the recent focus on the length of the lumbar spine in hominin evolution (Lovejoy and McCollum, 2010; O’Neill et al., 2018; Williams and Russo, 2015), the results of our study provide some evidence that the orientation of the trunk is just one factor, among many, that could impact 3-D gait kinematics. However, chimpanzees have a greater relative trunk size and a higher center of mass location of their trunk. Therefore, future studies could also modify the mass distribution of the trunk in humans to resemble the mass distribution of chimpanzees to quantify how the combined effects of mass distribution and anterior trunk flexion affect gait mechanics.

### Ground reaction forces

Consistent with the kinematic results, the GRFs were more chimpanzee-like in the human crouched posture conditions than the normal human gait condition. The GRFs in the crouched posture conditions are consistent with previous studies (Aminiaghdam et al., 2017; Grasso et al., 2000; Saha et al., 2008), with a greater first peak of the vertical GRF than second peak for the human crouched postures, and a less prominent trough during mid-stance for crouched postures than for normal walking. Despite this, the GRF patterns in each of the human crouched posture conditions were still quite distinct from the GRF patterns in bipedal chimpanzees, who exhibit a monophasic vertical GRF peak along with a smaller propulsive AP GRF when compared to human walking (Kimura et al., 1977; O’Neill et al., in review). These data indicate that crouched posture walking is insufficient to account for differences in vertical and AP GRF or patterns between human and bipedal chimpanzees walking, in contrast to some earlier inferences (Li et al., 1996). The differences in vertical GRFs between the crouched posture human gait and the chimpanzees give an indication that the COM motion is also different between these two gaits, which may impact the mechanical work performed on the center of mass during walking with implications for interspecific differences in walking economy (Demes et al., 2015; Kuo, 2007; Ortega and Farley, 2005; Wang et al., 2003)

In human gait, the second vertical GRF peak and propulsive AP GRF peak are dominated by ankle plantar flexor muscles (e.g. Liu et al., 2008; Winter, 1983). In bipedal chimpanzee walking, there is a greater reliance on the hip than the ankle in the second-double support period of the stance phase (O’Neill et al., in review; see also Sockol et al., 2007). The fact that humans retain a distinct push-off in crouched walking may reflect fundamental differences in relevant musculoskeletal traits of the human ankle (e.g., enlarged soleus; O’Neill et al., 2013) and foot (e.g., greater midfoot mobility; Holowka et al., 2017) as compared to bipedal chimpanzees (O’Neill et al., in review). Moreover, the vertical and AP push-off forces in human crouched posture walking fall between normal human walking and bipedal chimpanzee walking (Fig. 5), which combined with a distinct heel-strike in human crouched posture walking due to a dorsiflexed ankle angle (Fig. 4), suggest caution when inferring the habitual walking posture (upright or crouched) of extinct hominins from footprints, such as those preserved at Laetoli (e.g., Crompton et al., 2012; Hatala et al., 2016; Raichlen et al., 2010). Overall, these data show that human crouched posture gait remains distinct from chimpanzee gait in several important domains (pelvis motion, ankle angles, and GRFs) in ways that are likely critical to the interpretation of the evolution of hominin bipedalism.

### Limitations

One potential limitation of this study was the human subjects practiced each of the crouched posture conditions for only several minutes, rather than using the gait patterns for extended periods. A relatively short practice time was used in part because humans can readily walk with crouched postures, but the task can lead to muscle fatigue (e.g., Carey and Crompton, 2005) and we did not want our subjects to become fatigued during the testing session. Since the crouched posture conditions were not common tasks for the human subjects, it is possible that some other small modifications in gait mechanics would occur with a longer practice session or adaptation period (Sánchez et al., 2021; Selgrade et al., 2017). Additionally, this study built upon previous work by including instructions to modify human gait towards that of a chimpanzee in both the sagittal and frontal planes. However, it is possible that different types of feedback or instruction could be provided to the subjects to guide them towards an even more chimpanzee-like gait than what was observed in our data. However, additional instructions could become overwhelming to subjects as it would likely approach the limits of their working memory capacity to handle several instructions at once (Buszard et al., 2017). Future studies could implement multiple practice sessions or real-time visual feedback to further modify specific features of the kinematics or ground reaction forces in human crouched posture gait. Furthermore, utilizing a dimensionality reduction method, such as principle components, to give low-dimensional feedback rather than a set of multiple instructions (Day and Bastian, 2019; Day et al., 2019) could also allow for a closer match between human crouched posture and chimpanzee gait. Still, the series of instructions given in our study represents the most comprehensive step to date at instructing and measuring humans walking with a chimpanzee-like gait, and therefore represents an important advancement in directly comparing human and chimpanzee crouched posture gait mechanics.

### Conclusion

Human crouched posture conditions produced a more chimpanzee-like gait than normal human gait in pelvis list, hip and knee flexion-extension. However, substantial interspecific differences were evident in pelvis transverse plane rotation, hip abduction-adduction and mediolateral rotation, ankle plantar flexion-dorsiflexion and 3-D GRFs. These results suggest that human crouched posture walking captures a rather limited subset of the characteristics of 3-D kinematics and ground reaction forces, even with more elaborate instructions than is typical for human studies of this type. As such, human crouched walking is an inadequate proxy for chimpanzee or facultative bipedalism, despite what has previously been implied (e.g., Carey and Crompton, 2005; Li et al., 1996; Wang et al., 2003). It appears that constraints imposed by human anatomical structure cannot be overcome through guided instruction alone. Indeed, crouched posture walking in humans remains distinct from bipedal chimpanzees in numerous ways, suggesting some general caution when using human gait to understand the locomotor patterns of hominin species with distinct musculoskeletal designs (e.g., Carey and Crompton, 2005; Crompton et al., 1998; Raichlen et al., 2010). However, understanding why differences persist, even when attempting to recreate chimpanzee-like gait mechanics, can lead to novel insights linking musculoskeletal structure with gait mechanics, such as the proposed link between frontal plane knee angulation and hip abduction-adduction patterns during gait. In addition to musculoskeletal morphology and posture, other factors also likely play important roles in determining gait mechanics in humans and non-human primates such as the neural control of motion, metabolic energy expenditure, or dynamic stability and balance. Human crouched posture experimental studies may be most fruitful when used to test general principles of human musculoskeletal function, such as motor control (e.g., Grasso et al., 2000) or determinates of the metabolic costs of walking (e.g., Foster et al., 2013). Altogether, this study provides a comprehensive data set to illustrate the gait mechanics used by humans and chimpanzees during crouched posture bipedal gait, and the remaining differences between these species demonstrates how the distinct morphology of each species can impact bipedal gait.

## Acknowledgements

The authors thank R. Wedge for assistance with the human data collection. Thanks to K. Fuehrer for animal care and training.

## Competing Interests

No competing interests declared.

## Funding

This study was supported by the National Science Foundation (NSF) awards BCS 0935327 and BCS 0935321. Additional support was provided by a University of Massachusetts Amherst dissertation research grant.

## Data Availability

Average human and chimpanzee kinematic and ground reaction force data will be made available at https://simtk.org/projects/chimphindlimb upon publication.

## Notes

### Competing Interest Statement

The authors have declared no competing interest.

